# VIACOT: a web server for interactive visualization and comparison of cell lineage trees

**DOI:** 10.1101/2024.11.22.624819

**Authors:** Junnan Yang, Zizhang Li, Xiaoyu Zhang, Feng Chen, Peng Wu, Wenjing Yang, Xingxing He, Yuyan Shan, Jian-Rong Yang

**Affiliations:** Department of Biomedical Informatics, Zhongshan School of Medicine, Sun Yat-sen University, Guangzhou 510080, China; Department of Genetics and Cell Biology, Zhongshan School of Medicine, Sun Yat-sen University, Guangzhou 510080, China; Key Laboratory of Tropical Disease Control, Ministry of Education, Sun Yat-sen University, Guangzhou 510080, China

## Abstract

The cell lineage tree (CLT) records some or all cell division events during the development of multicellular organisms, as well as the functional characteristics of terminal cells, and is therefore commonly considered one of the most fundamental types of biological data. The combining of single-cell RNA sequencing with DNA barcoding has recently led to the rapid accumulation of CLT datasets. Unlike biological sequence data, web-based analysis and visualization tools for CLT data are currently lacking. In order to facilitate studies surrounding CLT, we built a webserver named VIACOT (Visualize Interactively And COmpare Trees). VIACOT is not only a comprehensive repository of published CLT datasets, but also a web tool that enables three preliminary analyses of preprocessed CLT datasets, including. (i) interactive visualization of CLT allowing fine-grain or coarse-grain exploration. (ii) calculations of correlations among single-cell metrics, such as gene expression levels, either using the original metrics or their phylogenetic independent contrasts. (iii) comparison among different CLTs to identify differences in cell fate. We anticipate that VIACOT will be a useful tool for exploration and biological interpretation of CLTs.

## Introduction

Each multicellular organism begins as a single zygote, which undergoes multiple rounds of division to give rise to a large number of diverse cells. The developmental cell lineage tree, in which the root represents the zygote and the branching events represent cell divisions, records the entire developmental process from a cellular perspective (1,2). On a broader definition, cell lineage tree can also be the record of continuous cell divisions starting from any dividable cell (3). First fully resolved developmental cell lineage tree was reported in 1983 by John Sulston and colleagues for *Caenorhabditis elegans* (2). This lineage tree has greatly aided our understanding of animal development, and has been honored by the 2002 Nobel Prize in Physiology or Medicine (4).

The resolution of a lineage tree by microscopic imaging, as performed by Sulston and colleagues in *C. elegans (2)*, is dependent upon the transparency of the body, the limited number of cells, and the deterministic development of the animal. These criteria are largely violated in more complex animals, such as humans and mice. Recent combination of DNA barcode (5-8) and single-cell transcriptomics has enabled lineage resolution for these more complex animals (9-21). For example, Raj and colleagues developed scGESTALT to trace the cell lineage tree of zebrafish brain (16). Unlike image-based lineage trees, in which all cells within the organism are connected by a binary tree, the new method typically generates cell lineage trees with polytomy (because some division events are not resolved), does not capture all cells within the organism (only a small percentage of all cells are recovered by single-cell RNA-seq (22)), and does not contain information about branch lengths (but see (23)). Limitations aside, this new type of lineage tree data typically contains hundreds of branches and terminal cells (tips on the tree). In order to efficiently understand the data, new specialized tools are required to visualize the complex data of the lineage trees.

To aid studies surrounding lineage trees, we hereby developed VIACOT (Visualize Interactively And COmpare Trees), an easy-to-use, open-source web server created specifically for the interactive visualization and comparison of cell lineage trees. VIACOT requires only topology of the tree such that it is compatible with DNA-barcode-based lineage trees. Furthermore, 29 public lineage trees obtained via imaging or DNA-barcoding have been incorporated into the web server. VIACOT is readily usable on any personal computer equipped with a mainstream modern browser without the installation of additional software. We expect that VIACOT will facilitate the close examination and comparative analysis of cell lineage trees by developmental biologists and evolutionary biologists interested in the phenotypes related to cell lineage trees.

## Results

### Overview of VIACOT

VIACOT is built as a JavaWeb server that uses the Spring Boot framework. We utilized vue.js, elementUI, and ApacheECharts (24) for interactive tree visualization, MySQL and MyBatis for storage and query of lineage tree data. VIACOT does not require any additional software to be downloaded or installed, other than a modern browser, which is already present on most personal computers. The browsers and operating systems listed in Table 1 have been tested and found to be compatible with VIACOT.

**Table 1.** Operating systems and browsers tested for compatibility.

| OS | Version | Firefox | Chrome | Safari |
| --- | --- | --- | --- | --- |
| Linux | CentOS 8.0 | 97.2.0 | 98.0.4758.102 | / |
| Windows | 10 | 97.2.0 | 98.0.4758.102 | / |
| MacOS | 10.16.1 | 97.2.0 | 98.0.4758.102 | 15.1.2 |

The VIACOT website has five major pages: Home, Search, Visualization, Download, and About Us, all easily accessible via the top navigation bar (Fig. 1A), and is provided in both English and Chinese (Fig. 1B). The Home page contains a brief introduction to VIACOT. Users can search VIACOT lineage tree datasets by specie names on the Search page. And the search can be further refined by limiting the number of cells within the tree so that only trees with a desired complexity are displayed. By clicking the “view” link to the right of each dataset, the user is redirected to the Visualization and Comparison page, with the corresponding tree loaded and displayed according to the default settings. The Visualization page allows users to view the cell lineage tree and interact with it by expanding or collapsing individual subtrees. Users can view just one tree in the single tree mode, or compare two genes within the same tree in the expression comparison mode, or compare two lineage trees in the tree comparison mode. All processed datasets in VIACOT are available for download in a structured format from the Download page. Last but not least, users may submit suggestions or new tree datasets to VIACOT by using the contact information listed on the About Us page.

**Fig. 1.**
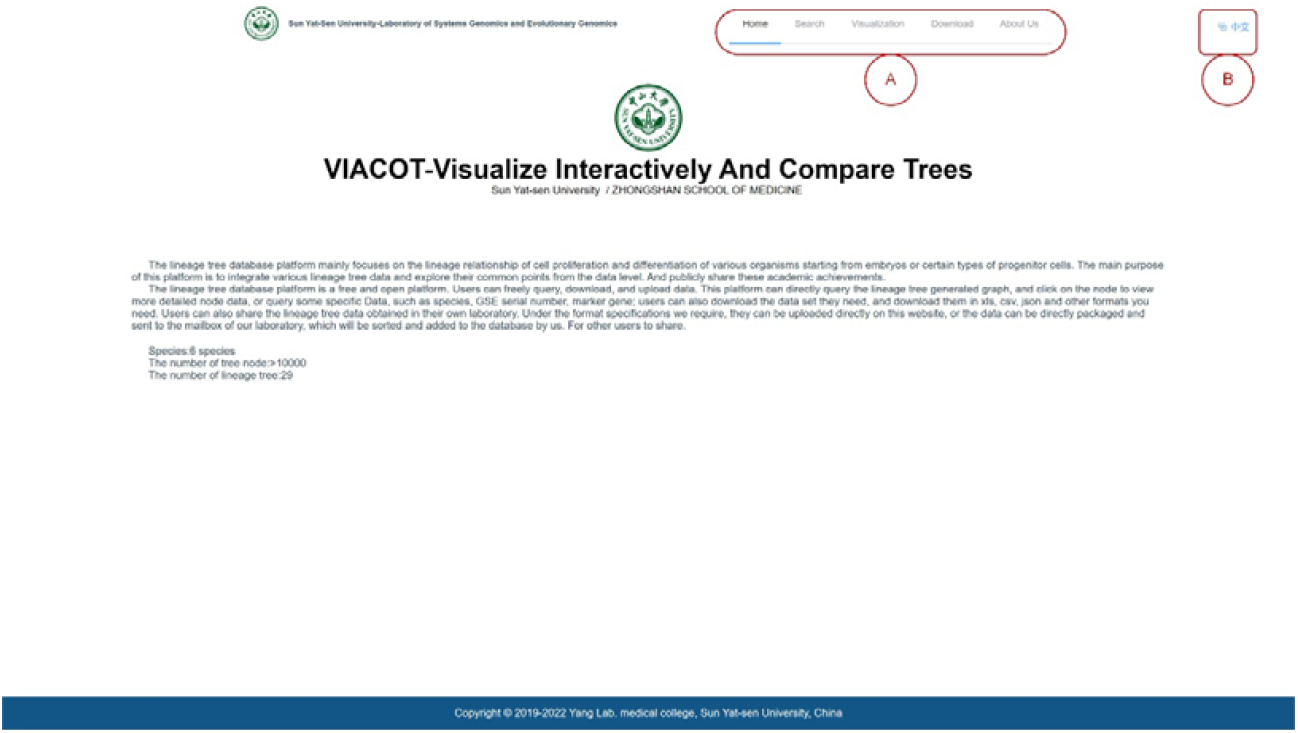
Overview of VIACOT. **(A)** Navigation bars to the five major pages of VIACOT: Home, Search, Visualization and Comparison, Download, and About Us. **(B)** Button to switch language between Simplified Chinese and English.

### The lineage tree datasets

A collection of pre-processed datasets of cell lineage trees was provided with VIACOT (**Table 2**). For these trees, interactive visualization that allows collapse and expansion of subtrees (as facilitated by VIACOT) is particularly useful, as they usually contained a large number of cells, which makes visualization by static image extremely challenging.

**Table 2.**
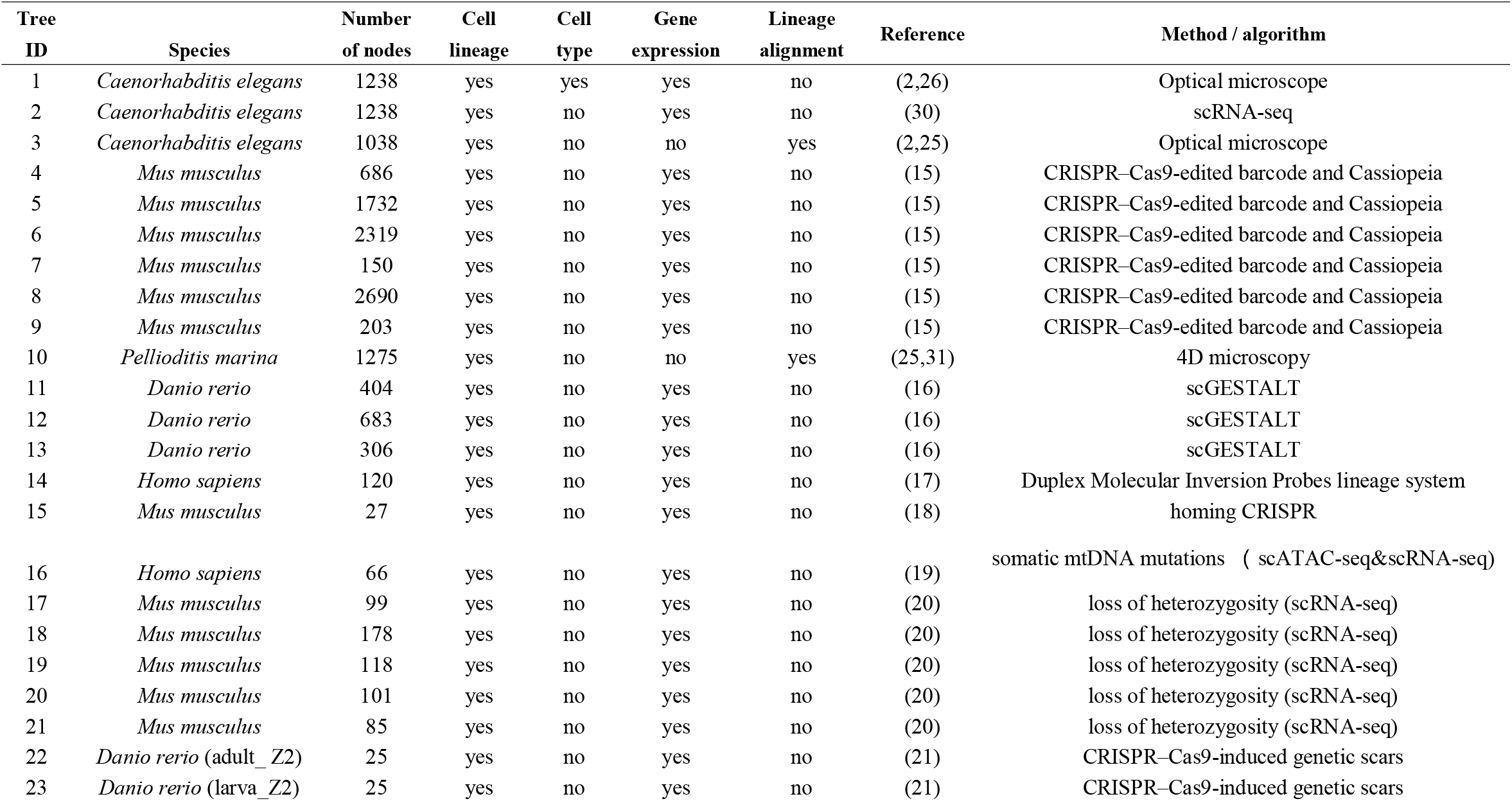
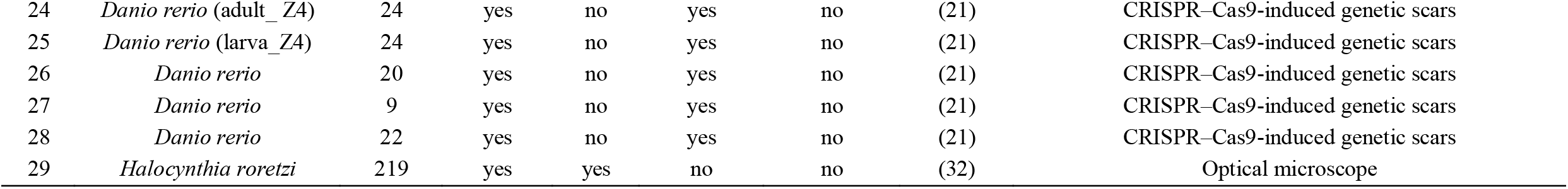
List of lineage tree datasets included in VIACOT.

On the Search page, users can search this collection of lineage trees. The full collection is displayed by default, or it can be reached by resetting the search criteria (Fig. 2C). On top of the Search page, there are two search criteria that may be used independently or in combination for each search (Fig. 2). On the one hand, species names may be entered for the search (Fig. 2A). On the other hand, the range of cell numbers within the tree can be selected (Fig.2B). Following each search, the list of trees matching the search criteria will be displayed at the bottom half of the Search page as a table, where each tree will be shown as a row (Fig.2D). Six columns are included in the table, including the tree name, species name, experimental or algorithmic method used to obtain the tree, the number of nodes within the tree, the publication describing the tree, and a link to the interactive visualization of the tree.

**Fig. 2.**
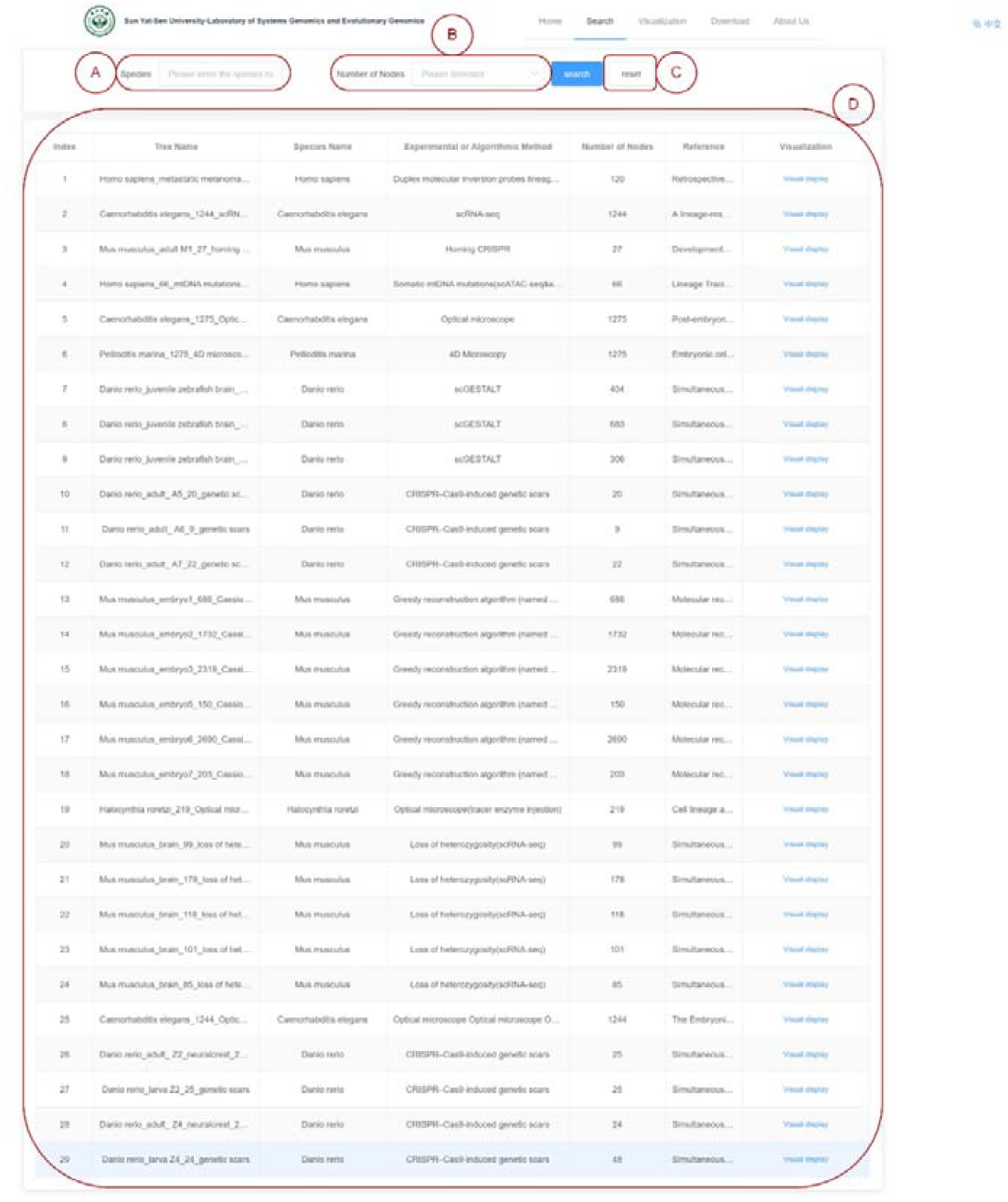
The Search page of VIACOT and the list of stored lineage trees. (**A-C**) Users can input species name **(A)** or select the range of nodes in the tree **(B)** as the search criteria, or press the reset button **(C)** to clear the search and list all available trees. **(D)** The table listing the search results. See also Table 2.

### Interactive visualization of cell lineage trees

The Visualization page offers three viewing modes, namely the single tree mode, the expression comparison mode and the tree comparison mode. The single tree mode displays a cell lineage tree (Fig. 3A) as a cladogram. The tree is not necessarily bifurcating, but may be multifurcating. By default, the tree is displayed horizontally (with the root on the left and the branches going rightwards), but the user can change the direction to vertical (with the root on top and the branches on the downwards), or radial (with the root at the center and the branches radiating outwards) via the “direction” option (Fig. 3B), or reset to the default display with the “reset” button (Fig. 3C).

**Fig. 3.**
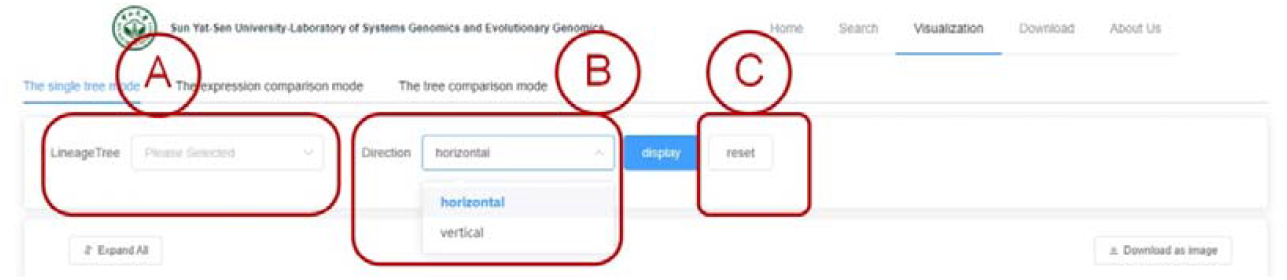
Basic options in the single tree mode of visualization. **(A)** Select the one tree to display in the single tree mode. **(B)** Choose the direction in which the tree will be displayed. It can be vertical (root at the top, branches downward) or horizontal (root at the left, branches rightward). **(C)** Reset the tree to the default view.

For any specific tree, it will be initially displayed at a depth (number of internal nodes since, including, the root) of two (Fig. 4). There are multiple ways to interact with the tree. There is an “expand all” button (Fig. 4A) that can be used to expand the whole tree. By left-clicking a node, it will be expanded and all its daughter nodes will be displayed unless it is a leaf node. A right-click on a node will display the subtree rooted at the clicked node, allowing the user to “zoom-in” and explore the subtree. Upon hovering the mouse pointer over a node, a message box (Fig. 4B) will appear showing the cell id, depth, tree name, and cell type information (if cell type data is present) for that node. By clicking on the “reset” button, you can return to the initial view of the full tree. Lastly, the user can download an image of the tree as seen by clicking the “download as image” button (Fig. 4C).

**Fig. 4.**
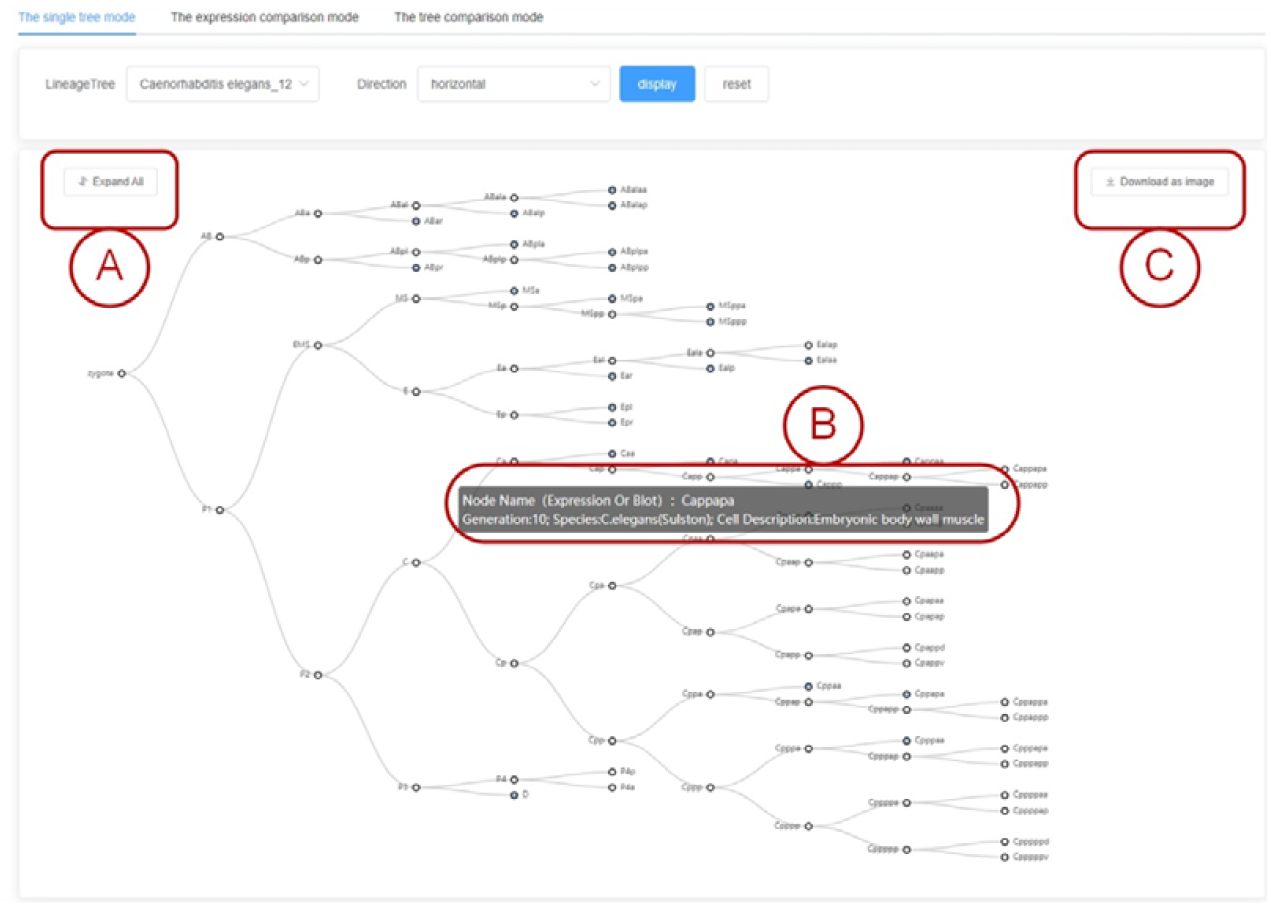
Interactive visualization of a single tree. **(A)** Click on the button to expand all internal nodes and display the entire tree. **(B)** An information box containing details of the node over which the mouse cursor is hovering. **(C)** Click the button to download the current tree as an image, with individual nodes expanded or collapsed as displayed on the screen.

Additionally, in the two comparison modes (expression comparison mode and the tree comparison mode), two lineage trees are displayed face-to-face, and they can be interacted with similarly to the single tree mode (except that the “zoom-in” by right-clicking a node is not implemented therein). These comparison modes serve two primary purposes. For the expression comparison mode, the expression of different genes can be compared for the cells within the same tree. For the tree comparison mode, comparing different lineage trees is also made possible with the tree alignment result by the DELTA algorithm (25) (see below). In the following sections, we will discuss in more detail these two main functions of the comparative mode.

### Visualizing and comparing gene expression on lineage trees

For scRNA-seq base cell lineage trees, single-cell transcriptomes, i.e. the mRNA levels of genes within each terminal cell, are available. Similarly, the protein expression of some genes has been determined by time-lapse imaging in *C. elegans* (26). Gene expression closely reflects cellular states, and is therefore an indispensable element in the biological interpretation of the lineage tree. In the expression comparison mode of Visualization, the terminal nodes of the tree are colored based on whether a specific gene is expressed (Fig. 5A), enabling the investigation of gene expression across the lineage. Various patterns of clustering (Fig. 5B) or dispersion (Fig. 5C) of gene expression can then be observed in the tree.

**Fig. 5.**
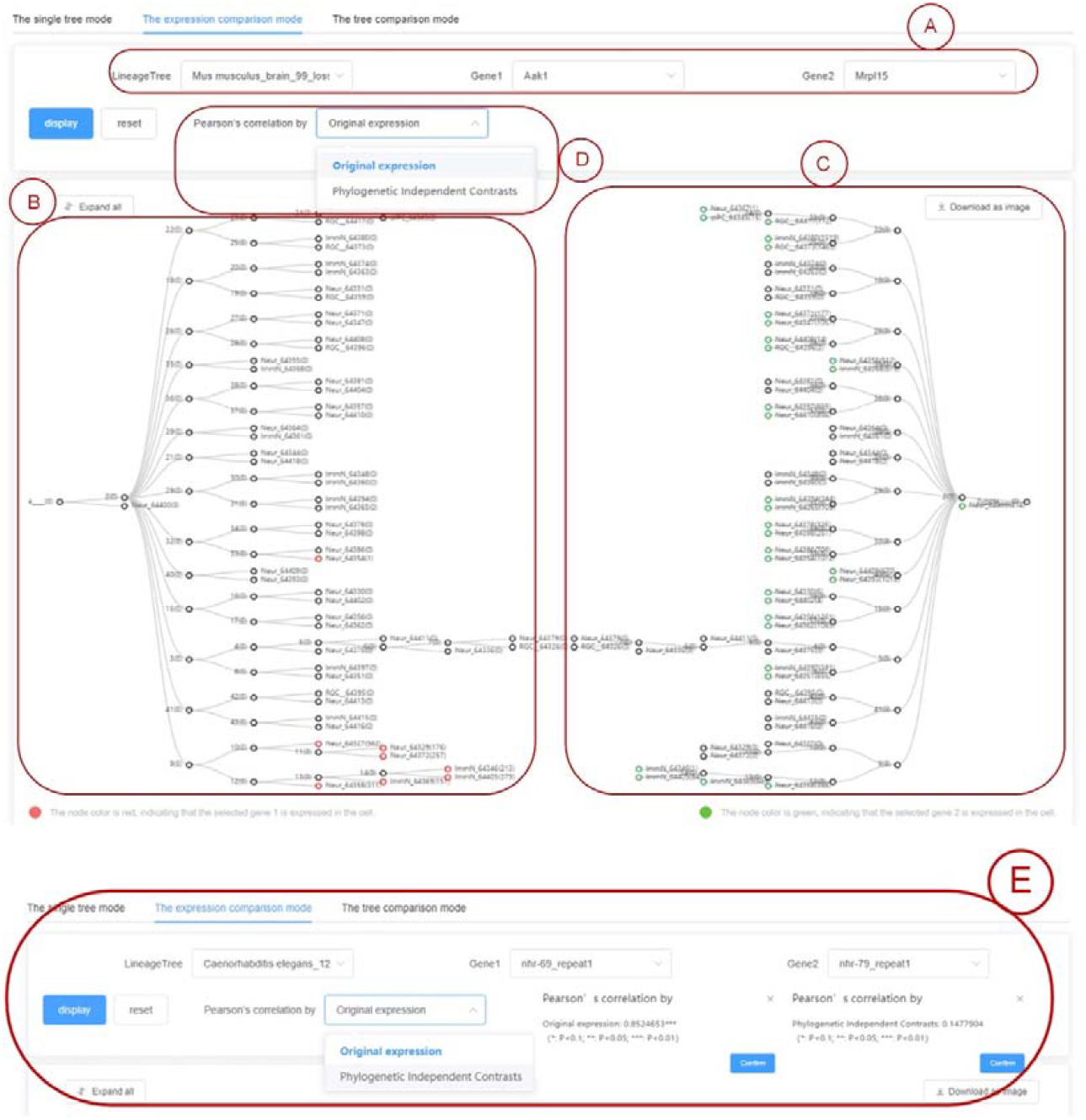
Expression comparison mode of visualization. **(A)** The lineage tree and the two genes whose expression will be shown should be selected. **(B-C)** Examples of a gene with clustered expression (B) and another gene with dispersed expression (C) on the lineage tree are presented. **(D)** The Pearson’s correlation coefficient can be calculated either using the original expression levels or using the Phylogenetic Independent Contrasts (PICs) of the two genes. **(E)** A pair of genes with significant correlation when using the original expression levels but not PICs.

Displaying gene expression levels in the comparison mode as node colors facilitates further comparisons between genes within the context of the lineage tree. The inclusion of the lineage tree is particularly important in this regard, since shared ancestry (as reflected in the lineage tree) creates statistical interdependence among cells, which can impair the comparison, for example, by creating false correlations (27). To shed light on related analyses, VIACOT calculates the Pearson’s correlation between the two selected genes by using the original and the Phylogenetic Independent Contrasts (27) of the expression level (Fig. 5D). A pair of genes that displays significant correlation using the original expression but not PIC is shown as an example to illustrate how false correlations created by shared ancestry of cells can be controlled by PIC (Fig. 5E).

### Comparison of different trees facilitated by DELTA

We have previously developed an algorithm named DELTA (developmental cell lineage tree alignment) to compare different cell lineage trees. By exhaustively searching for optimal correspondence between individual cells from different trees while retaining their topological relationships, DELTA quantifies phenotypic similarity between two lineage trees, or in other words, aligns them(25). We have demonstrated that aligning lineage trees, similar to aligning sequences, can assist with functional and evolutionary inferences regarding genetic and developmental programs and cell fates(25). For the purpose of facilitating these DELTA-based analyses of lineage tree alignments, the tree comparison mode under Visualization allows users to view the DELTA alignment results. Due to the subjective definition of cell types and the limited resolution of cell phylogeny, biologically meaningful alignments are not yet available for the scRNA-seq-based lineage tree. Accordingly, we presented the global alignment between *C. elegans* and *P. marina* as an example in VIACOT (Fig. 6A).

**Fig. 6.**
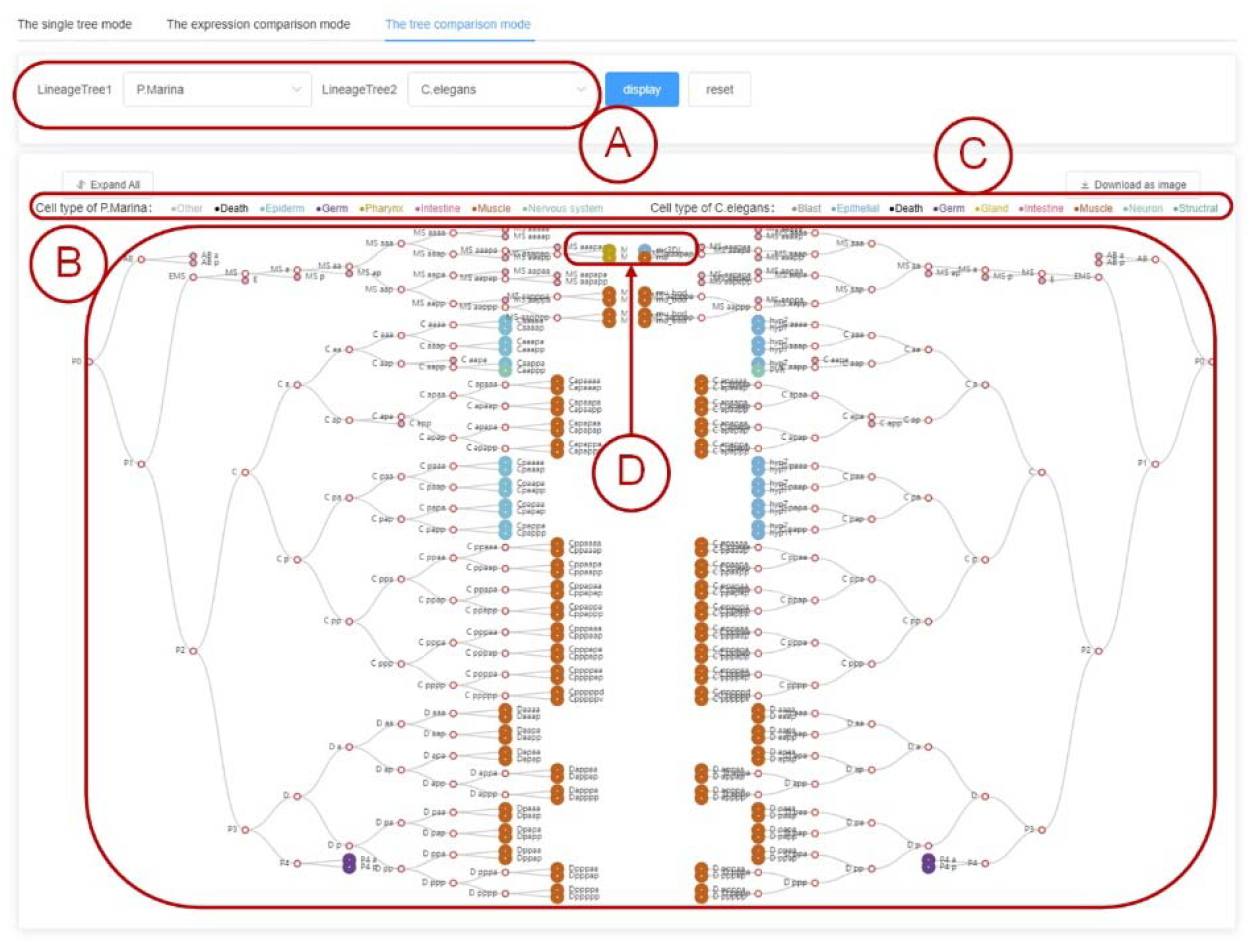
Comparative visualization of two different lineage trees. **(A)** Select the two lineage trees that will be aligned and displayed. **(B)** The two lineage trees will be displayed face-to-face, with aligned nodes expanding and collapsing in synchrony. **(C)** The colors of the nodes indicate the terminal cell types of the lineage trees. **(D)** The evolutionary changes of cell fates highlighted by aligned nodes with different colors.

When using the tree comparison mode, the two lineage trees are synchronized such that expanding or collapsing any node in one of the two aligned trees will have the same effect on the aligned node in the other tree (Fig.6B). Also, the functional type of the terminal cells is indicated by the color of the node (Fig.6C). Therefore, the cell types of aligned terminal cells can be visually compared, allowing evolutionary changes of cell fates to be highlighted by contrasting colors (Fig. 6D).

## Discussion

### Comparison with other databases and tools

VIACOT is, to our knowledge, the first web server built for interactive visualization of cell lineage tree that allows node annotation (gene expression or cell type) and tree comparison. With more and better scRNA-seq-based lineage tree data becoming available, we envision that VIACOT will greatly help biological interpretation of lineage tree data.

To date, there have been a limited number of web-based tools for analyzing cell lineage tree data, including MorphoNet(28) and CeLaVi(29). There are, however, significant differences between VIACOT and these two tools. As suggested by the name, MorphoNet’s primary function is to visualize morphological information relating to three-dimensional objects, which range from cells to tissues to organs. Visualization of cell lineage is only a minor component of MorphoNet’s functionality, as there are only a small number of ways that users can interact with cell lineage, and it is not possible to display lineage trees with many cells. The process of installing, registering, and obtaining access to data further hampers MorphoNet’s ease of use. CeLaVi is a tool that integrates four example datasets of cell lineages from two species, as well as spatial information of individual cells. However, CeLaVi allows neither comparative visualization of two trees, nor collapsing/expanding of non-bifuricating nodes. More importantly, CeLaVi is incompatible with gene expression data, which is a unique feature of the ever-increasing scRNA-seq-based data. In contrast, VIACOT allows interactive visualization of more than twenty prestored lineage datasets, displays the expression of specific genes as colors for the terminal nodes of the cell lineage tree, permits simultaneous visualization of two distinct lineage trees in an aligned manner, as generated by DELTA.

### Limitations and perspectives

We have collected major cell lineage datasets published before December 2021. More datasets will undoubtedly appear in the future. It is likely that there will be a need to visualize custom lineage tree datasets besides those already available in VIACOT. Nonetheless, we do not permit data upload by users due to the lack of a standard data format, as well as the complexity of lineage tree data, which makes automatic processing of such data difficult. Users are, however, invited to submit their data to us for processing and uploading to VIACOT. In the future, we intend to add a feature that will let users upload and view their own lineage tree data.

Furthermore, VIACOT performs most of its computations on the server side. Thus, good connectivity, speed and bandwidth of the network, a short response time, and a large memory on the server are required. To further improve VIACOT, we plan to use web caching technology and optimize the allocation of storage space and the global index of the database in order to improve the loading speed of lineage trees.

We anticipate that VIACOT, with its continuous enhancements, will facilitate the close examination and comparative analysis of cell lineage trees, an important phenotype which is becoming increasingly available.

## Data availability

VIACOT is accessible on http://49.234.48.196. All datasets in VIACOT, as listed in Table 2, can be downloaded from the download page.

## Code availability

VIACOT is completely open-source on github (https://github.com/JunnanYang2022/VIACOT-server)

## Acknowledgment

This work was supported by the National Key R&D Program of China (grant number 2021YFF1200904 to J.-R. Y.) and the National Natural Science Foundation of China (31871320 and 32122022 to J.-R. Y.).

## Author Contribution Statement

J.-R.Y. conceived the idea, designed and supervised the study. J.Y., Z.L., X.Z., F.C., P.W., W.Y., Y.S. and J.-R.Y. acquired and analyzed data. J.-R.Y. contributed new analytical tools and acquired funding. J.Y. and J.-R.Y. designed and developed the software. J.-R.Y. and J.Y. wrote the paper with input from all authors.

